# Associative learning in the protozoan Stentor coeruleus

**DOI:** 10.64898/2026.02.25.708045

**Authors:** Nhi Doan, Austen Theroux, Tejas Ramdas, Samuel J. Gershman

## Abstract

The capacity for associative learning in protozoa has been a matter of longstanding controversy. In a series of Pavlovian conditioning experiments with the ciliate *Stentor coeruleus*, we show that temporally pairing weak and strong mechanical stimuli results in a transiently enhanced contraction response to the weak stimulus. Control experiments rule out several alternative explanations, such as non-associative sensitization or arousal. Parametric manipulation of the conditioning protocol’s temporal structure revealed a systematic dependence of learning on the inter-trial and inter-stimulus intervals, though not in the form classically observed in animals. A simple mathematical model, combining associative learning with habituation, can explain why enhancement is transient, and accurately fits the learning curve at the aggregate level. We conclude that *Stentor coeruleus* appears capable of associative learning, suggesting an ancient evolutionary origin that preceded the emergence of multicellular nervous systems.

## 1 Introduction

When one event or stimulus reliably predicts another, many animals acquire an anticipatory response that reflects their learned expectations. The most famous demonstration is Pavlovian conditioning ^1^, where an initially neutral stimulus (the conditioned stimulus, or CS) is predictive of an appetitive or aversive stimulus (the unconditioned stimulus, or US). This procedure results in the animal’s production of a conditioned response (CR) to the CS. For example, a rat that reliably hears a tone CS several seconds before a footshock US will begin to freeze (the CR) in response to the tone.

Many theories posit that Pavlovian conditioning arises from the formation of associative links between stimuli ^2^, though this hypothesis is still a matter of controversy ^3,4^. So ingrained is this way of thinking that Pavlovian conditioning, and related phenomena such as operant conditioning, are referred to collectively as *associative learning*. The mechanism underlying Pavlovian conditioning (and associative learning more generally) is widely believed to be the Hebbian modification of plastic synapses ^5,6^, though this is also a matter of controversy ^7,8^. According to this hypothesis, different neurons correspond to the CS, US, and CR; learning corresponds to the strengthening of the synaptic strength between neurons representing the CS and the US. This strengthening depends on the co-occurrence of firing between the CS and US neurons. The effect of learning is to activate the US neuron in anticipation of the US itself, thereby driving downstream behavioral responses (the CR).

Given the centrality of synaptic plasticity to this story, it is tempting to think that associative learning is an ability restricted to organisms with synapses. However, evidence exists that organisms without synapses (indeed without any brains at all), such as some protozoa, may be capable of associative learning ^9–12^. For example, Gelber ^13^ demonstrated that the ciliate *Paramecium aurelia* produced a Pavlovian approach behavior (the CR) towards a platinum wire (the CS) that had been previously coated with bacteria (the US). Gelber’s studies, and other demonstrations of Pavlovian conditioning in protozoa, have been disputed ^14–16^, such that we currently lack decisive evidence. If such evidence can be procured, it could suggest that associative learning is supported in more than one biological substrate, inviting a reassessment of the evolution of associative learning and of the relationship between synaptic and non-synaptic mechanisms of learning in more complex organisms.

Our goal is to close the evidential gap by studying a form of Pavlovian conditioning in the ciliate *Stentor coeruleus* (here *Stentor* for short), a large (1mm) protozoan that anchors to the bottom of ponds and filter feeds through its trumpet-like oral apparatus. The cognitive abilities of *Stentor* have fascinated investigators for over a century^17,18^, and its capacity for learning has recently come under renewed scrutiny^19,20^. Here we report what is, to the best of our knowledge, the first demonstration of associative learning in *Stentor*.

Our setup builds upon the mechanosensory habituation protocol pioneered by David Wood^22^ and recently revived by Deepa Rajan, Wallace Marshall and their collaborators^19^. In this protocol, *Stentor* are exposed to a series of periodic mechanical stimuli (taps), which are aversive to the cells and produce a defensive response (contraction; Fig. 1A,B). With repeated stimulation, the cells reduce their contraction probability—the basic habituation phenomenon. We introduce a novel variation of this protocol, where a weak tap (which at baseline produces a low contraction probability) reliably precedes a strong tap. This produces a qualitatively different pattern of responses: contraction probability initially increases and then decreases. We will argue that this is a manifestation of associative learning, reproduced by a simple computational model. Our findings are consistent with experimental data from animals using a similar protocol ^23^. In the Discussion, we address the implications for fundamental questions about the mechanisms and origins of learning.

**Figure 1:**
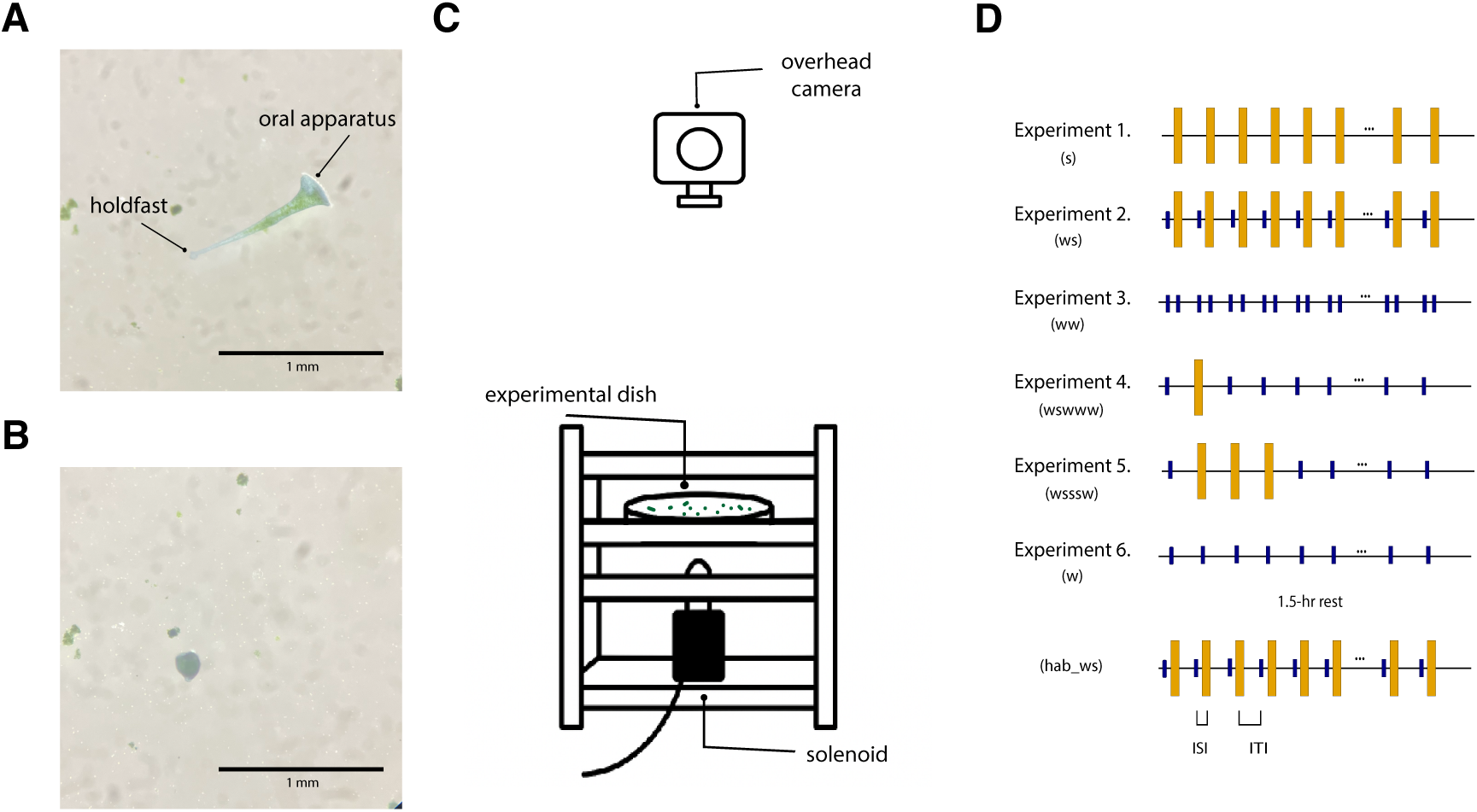
Studying Pavlovian conditioning in non-neural, single-celled *Stentor coeruleus*. **(A)** At rest, *Stentor* exhibits an elongated form with a holdfast attached to the substrate and a trumpet-like oral apparatus covered in cilia at the anterior end. **(B)** When exposed to mechanical stimuli, *Stentor* contracts its cortical myoneme, reducing body length to less than 50% of its original size within milliseconds ^21^. In their natural environment, mechanical cues can signal genuine threats (predators) or false alarms (water currents, falling debris). Fast contraction gives *Stentor* time and space to escape predators, but is also energetically expensive and prohibits feeding, creating pressure to distinguish threatening signals from benign stimuli. **(C)** The experimental setup comprises a custom-made device mounted on an isolation system to ensure *Stentor* receives no extra disturbance except timed mechanical stimuli. A camera and LED ring are positioned above the setup. The Petri dish containing *Stentor* is placed in the center of the box, right above the solenoid (black). When energized, the solenoid piston moves up, taps the dish, and falls back down. **(D)** Experimental protocols. The small bar indicates a weak tap; the large bar indicates a strong tap. ISI: inter-stimulus interval. ITI: inter-trial interval.

## 2 Results

### 2.1 *Stentor coeruleus* show associative learning

Stentor exhibit intensity-dependent habituation to mechanical stimuli^22^. We have reproduced this finding using a slightly modified version of the apparatus developed by Rajan et al. ^19^, shown in Fig. 1C. Our apparatus employs a solenoid to deliver timed mechanical taps, mounted on a vibration isolation platform. We verified stimulus delivery using both an independent audio monitoring system and an overhead camera. To prevent factors other than mechanical stimulation from influencing our results, we performed experiments across different times of day and with replicates from multiple independently cultured populations. More experimental details are provided in Methods.

We first replicated the high-force stimulation paradigm used in previous studies ^19,22^ (Fig. 1D, Experiment 1). Consistent with prior findings, *Stentor* exposed to repeated 60 high-force (strong) taps exhibited an initially high probability of contraction followed by a gradual decay (Fig. 2A), though we note that each underlying process at a single-cell level is likely much more abrupt^19^.

**Figure 2:**
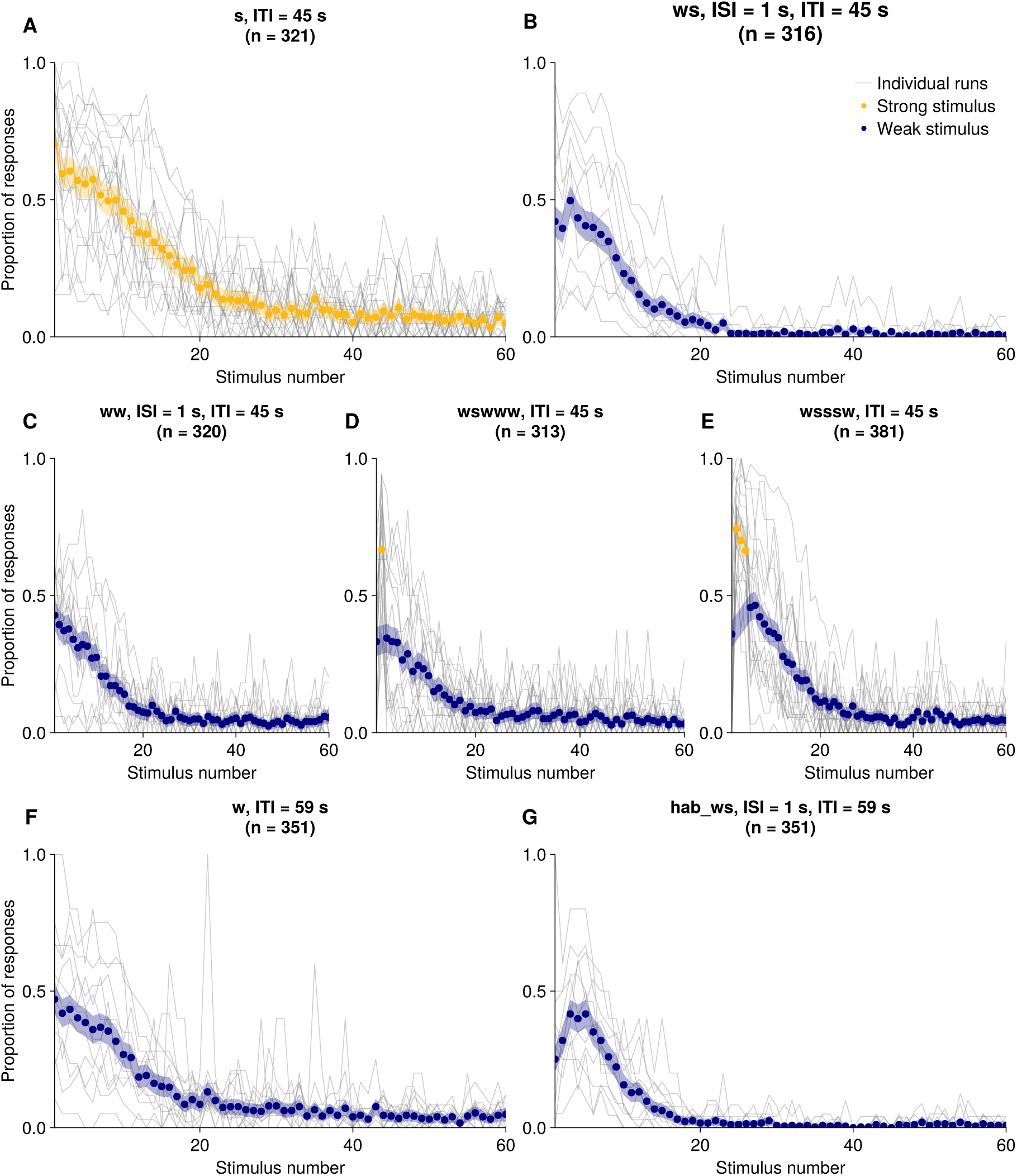
Stentor coeruleus exhibits associative learning that cannot be explained by nonassociative mechanisms. All protocols deliver 60 mechanical stimuli (45-s ITI, 1-s ISI unless noted). Navy/yellow points show mean weak/strong tap responses; gray lines are individual runs; shaded regions are 95% CIs, (mean ± 1.96 SEM; for proportions, 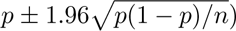. **(A)** Strong taps alone (n = 321 cells, 19 runs). **(B)** Weak-strong pairing (*n* = 316 cells, 10 runs). **(C)** Weak-weak pairing (*n* = 320 cells, 14 runs). **(D)** Single strong tap then weak taps (*n* = 313 cells, 11 runs). **(E)** Three strong taps then weak taps (*n* = 381 cells, 10 runs). **(F)** Weak taps alone (ITI=59s) (*n* = 351 cells, 10 runs). **(G)** After pre-habituation and 1.5-hr rest, weak-strong pairing produces robust non-monotonic responses despite *>*2-fold reduced baseline (*n* = 351 cells, same runs as **F**; ITI=59s).

We next developed a Pavlovian conditioning paradigm pairing strong and weak taps separated by a 1 second inter-stimulus interval (ISI). In this paired (“weak-strong” or ws) protocol, 60 weak taps reliably predicted incoming strong taps (Fig. 1D, Experiment 2), similar to Pavlovian conditioning where a CS predicts US. In their natural environment, this paradigm mimics small mechanical cues (such as minor water disturbances) that consistently precede genuine threats (such as approaching predators). If *Stentor* can learn this predictive relationship, cells should show enhanced responses to the initially weak stimulus as it acquires predictive value. We compared this to a “weak-weak” (ww) protocol with identical temporal structure but replacing strong taps with weak taps (Fig. 1D, Experiment 3). This manipulation preserves the temporal pairing structure while eliminating the adaptive incentive to learn: since weak taps alone pose minimal threat and elicit low baseline responses, there is no ecological benefit to predicting their arrival.

Cells in the weak-strong protocol exhibited a non-monotonic learning trajectory over the first ten trials (Fig. 2B), first increasing and then decreasing their contraction probability. This contrasted with the more typical habituation trajectory observed in the standard strong-only protocol (Fig. 2A) and weak-weak protocol (Fig. 2C). To test whether the weak-strong and weak-weak protocols elicit distinct response trajectories, we fit separate quadratic mixed-effects models for each condition using the first 10 stimuli. Each model predicted the number of contracted cells as a function of stimulus number and its quadratic term (stimulus^2^), with random intercepts across experimental runs to account for baseline run-to-run variability. The weak-strong condition exhibited both a significant negative quadratic coefficient (*β*_stimulus2_ = *−*0.144, *p <* 0.001) and a significant positive linear coefficient (*β*_stimulus_ = 0.934, *p* = 0.026), with the estimated peak at stimulus *−β*_1_*/*(2*β*_2_) *≈* 3.2. This reflects the non-monotonic response trajectory in which cells initially contracted but habituated over successive stimuli. Bayesian quadratic regression further supported this, yielding a posterior probability of 0.95 that the peak fell within the stimulus range. In contrast, the weak-weak condition showed no significant quadratic component (*β*_stimulus2_ = 0.014, *p* = 0.463) but a significant linear decline (*β*_stimulus_ = *−*0.531, *p* = 0.014). These patterns demonstrate that the two protocols produce qualitatively different responses over repeated stimulation.

To determine whether the elevation in responses seen in the weak-strong protocol reflected associative learning rather than non-associative sensitization, we conducted three control experiments. First, we tested whether a single strong tap causes non-specific arousal that could enhance subsequent responses (Fig. 1D, Experiment 4). We introduced a single strong tap following an initial weak tap, maintaining the same interval before delivering the next weak tap (Fig. 2D). If non-specific arousal drives response enhancement, we would expect elevated responses to the weak tap immediately following the strong tap. However, when comparing responses to the first weak tap (before strong tap exposure) and the third tap (the weak tap immediately after the strong tap), we found no significant difference (paired *t*-test, *p* = 0.94). This result rules out the possibility that response elevation is due to non-specific arousal following a single aversive stimulus.

Second, we examined whether multiple unpaired strong taps could induce non-specific sensitization sufficient to enhance subsequent responses (Fig. 1D, Experiment 5). This protocol was inspired by sensitization studies in *Aplysia*, which induce heightened arousal to a stimulus following repeated aversive events ^24^. If such non-specific sensitization drives the response enhancement observed in the weak-strong protocol, then exposure to multiple strong taps alone—even without predictive pairing—should be sufficient to elevate responses to weak taps. We exposed cells to three consecutive strong taps, then switched to weak taps while maintaining the same 45-s ITI (Fig. 2E). Comparing responses to weak taps after strong tap exposure with baseline responses to weak taps, we found no significant difference between the two population of responses (paired *t*-test, *p* = 0.23). This result suggests that cumulative exposure to aversive stimuli alone does not account for the response enhancement.

Third, we examined whether the response enhancement reflects learned prediction or simply sustained excitement to inherently salient weak taps. Unlike the ideally neutral CS in classical conditioning, weak taps in our protocol do elicit non-zero baseline responses. One alternative explanation is that the observed enhancement merely reflects cells maintaining this pre-existing responsiveness, with strong taps preventing normal habituation, rather than cells acquiring new anticipatory responses based on the predictive relationship.

To distinguish these possibilities, we reduced the initial salience of weak taps through pre-habituation. Cells received a series of weak taps (habituation phase), followed by a 1.5-hour rest period, then the standard weak-strong pairing protocol (Fig. 1D, Experiment 6). This pre-habituation successfully reduced initial responses: cells showed less than half the response probability to the first weak tap in the pairing phase compared to the standard protocol (Fig. 2G vs. B at stimulus 1). Critically, despite this two-fold reduction in baseline salience, pre-habituated cells exhibited more robust non-monotonic response patterns, with contraction increasing sharply during early pairing trials before declining (Fig. 2G, *β*_stimulus2_ = *−*0.242, *p <* 0.001). This result demonstrates that the predictive relationship between weak and strong taps drives the formation of new anticipatory responses, even when the CS has been rendered more neutral through pre-habituation.

In a further experiment, we varied the conditions of training and then tested under a common condition (Fig. 3A). To assess the robustness of our results, we ran two sets of conditions with different timing between training and testing and the length of pre-habituation periods, but each set of conditions follows the same structure: a pre-habituation period, a training period, and a test. The pre-habituation period here is designed to decrease initial responsiveness to the weak taps themselves, so that any subsequent change can be attributed to the difference in the training procedure. For training, we use sensitization and associative-learning protocols similar to those we previously reported, which are five weak-strong pairings (experimental, for short) and a weak tap followed by a series of strong taps (control; analogous to experiment 2 and 5 in Fig. 1). The experimental group received five weak–strong pairings, and the control group received a single weak tap followed by a series of strong taps. If the phenomenon we have observed reflects sensitization, then pairing should have no effect and the two groups should respond similarly at the common test. If the associative interpretation is correct, the two groups should differ: the experimental group should respond more to the weak tap, because for that group the weak tap has acquired predictive value through its pairing with the strong tap, whereas the control group should continue to habituate in the absence of such a predictive relationship.

**Figure 3:**
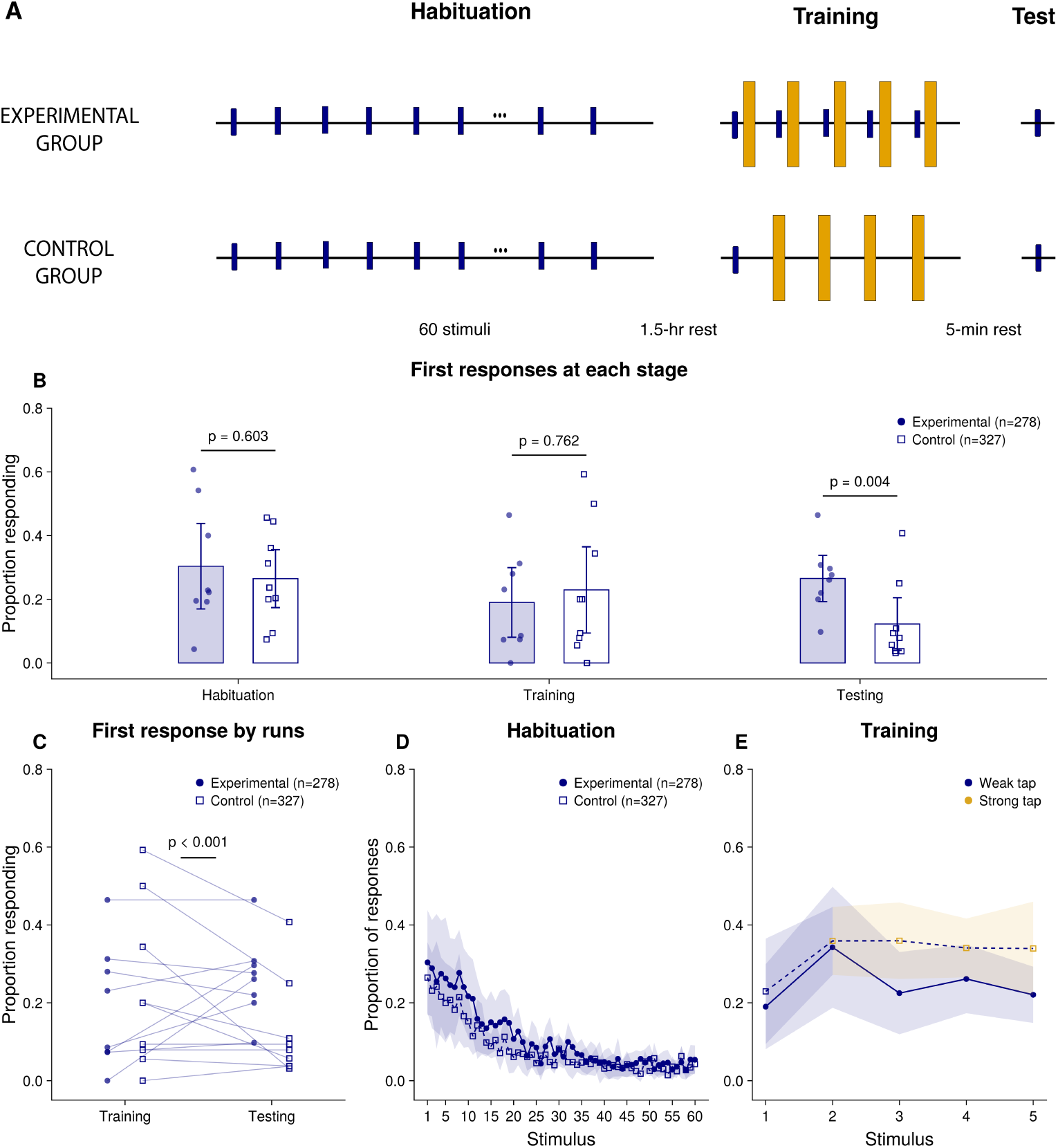
Training changes cells’ responses at a common test. **(A)** Experimental design for Set 1 data. Both groups receive identical pre-habituation (60 weak taps) and an identical test (one weak tap), and differ only during training: five weak–strong pairings (experimental) versus one weak tap followed by four strong taps (control). Navy lines/dots are weak taps; yellow ones are strong taps. Rests were 1.5 hr before training and 5 min before test. Set 2 (Fig. S2) follows the same design with different intervals. Filled navy circles represent experimental condition (*n* = 278, 8 runs); open squares are control (*n* = 327, 9 runs). **(B)** Responses to the first weak tap of each phase. Bars are means across runs, points are individual experimental runs, error bars are 95% CIs. Each *p* is for condition (response *∼* condition + (1*|*run), fit per phase). **(C)** The same first-tap responses at training and test, one line per run; *p* is the condition*×*phase interaction p-value. **(D)** Pre-habituation curve; points are means across runs, shading the 95% CI. **(E)** Training curve, as in (D). All models are mixed-effects logistic regressions (Bernoulli, logit link).

In Set 1 (experimental *n* = 278, control *n* = 327), the two groups responded similarly during pre-habituation (first stimulus p = 0.603, see Fig. 3B first column, D) and at the first stimulus of training (p = 0.762, Fig. 3B second column, E). At common test, after both groups went through the training period, two groups diverge significantly (condition*×*phase interaction *β* = *−*1.35, *p <* 0.0001, Fig. 3C). Specifically, the experimental group increased their responses to the weak tap from training to test, whereas the control group continued to decline, so that at test experimental cells were more likely to respond than controls (25.3% vs. 11.3%; *β* = 1.10, SE = 0.38, *z* = 2.85, *p* = 0.004; Fig. 3B last column). In Set 2 (experimental *n* = 348, control *n* = 173), training with paired weak–strong taps also elicited the same divergence (condition*×*phase interaction *β* = *−*1.57, SE = 0.37, *z* = *−*4.26, *p <* 0.0001, Fig. S2C), even though this set had different pre-habituation periods and rest times. Here the effect of condition reversed between the two phases: the control group responded more at the first stimulus of training (*β* = *−*0.67, SE = 0.35, *z* = *−*1.89, *p* = 0.059), while the experimental group responded more at test (*β* = 0.61, SE = 0.33, *z* = 1.84, *p* = 0.065, Fig. S2B second and last columns). If training did nothing, we would expect the two groups to have changed in the same way from training to test, regardless of their starting levels. Instead, the two groups changed in opposite directions in both data sets: the experimental group increased its responses while the control group became less responsive. This held in Set 1, where the groups started training at the same level, as well as in Set 2. We take this as evidence that the pairing between weak and strong taps drives the increase in responses, and that *Stentor* retain a memory of this association, which non-associative accounts like sensitization fail to explain.

Together, these results show that *Stentor* acquire anticipatory responses to weak stimuli based on their predictive relationship with strong taps. We confirmed that the increase in responses in our conditioning paradigm is not due to temporal structure, arousal following a single aversive stimulus, cumulative sensitization from repeated strong taps, or sustained excitement to inherently salient weak taps. With the common test procedure, we show *Stentor* maintain their memory beyond the training period.

### 2.2 Learning dynamics with varying inter-trial and inter-stimulus intervals

Having established that *Stentor* display associative learning, we next explored how temporal structures influence learning dynamics. In Pavlovian conditioning, both the inter-stimulus interval (ISI, delay between CS and US) and inter-trial interval (ITI, delay between successive pairings) determine how fast organisms acquire the learned responses ^25^. We therefore systematically varied ISI (1–20 s) and ITI (45–590 s) in Experiment 2 to test whether *Stentor* exhibits similar temporal sensitivity. We chose these ranges to allow full re-extension between contractions while keeping total experiment duration short. We observed non-monotonic response patterns in five of six conditions tested (Fig. 4A-E). At the longest timescales (ISI=10s, ITI=590s), the quadratic coefficient was positive (*β*_stimulus2_ = 0.061, *p* = 0.02), and the Bayesian analysis yielded a posterior probability of 0.0017 for a peak within the stimulus range, indicating that cells did not exhibit the inverted-U pattern observed in the other conditions (Fig. 4F). This boundary suggests that *Stentor* ’s temporal window for associative learning extends across seconds to minutes but not beyond 10-minute intervals.

**Figure 4:**
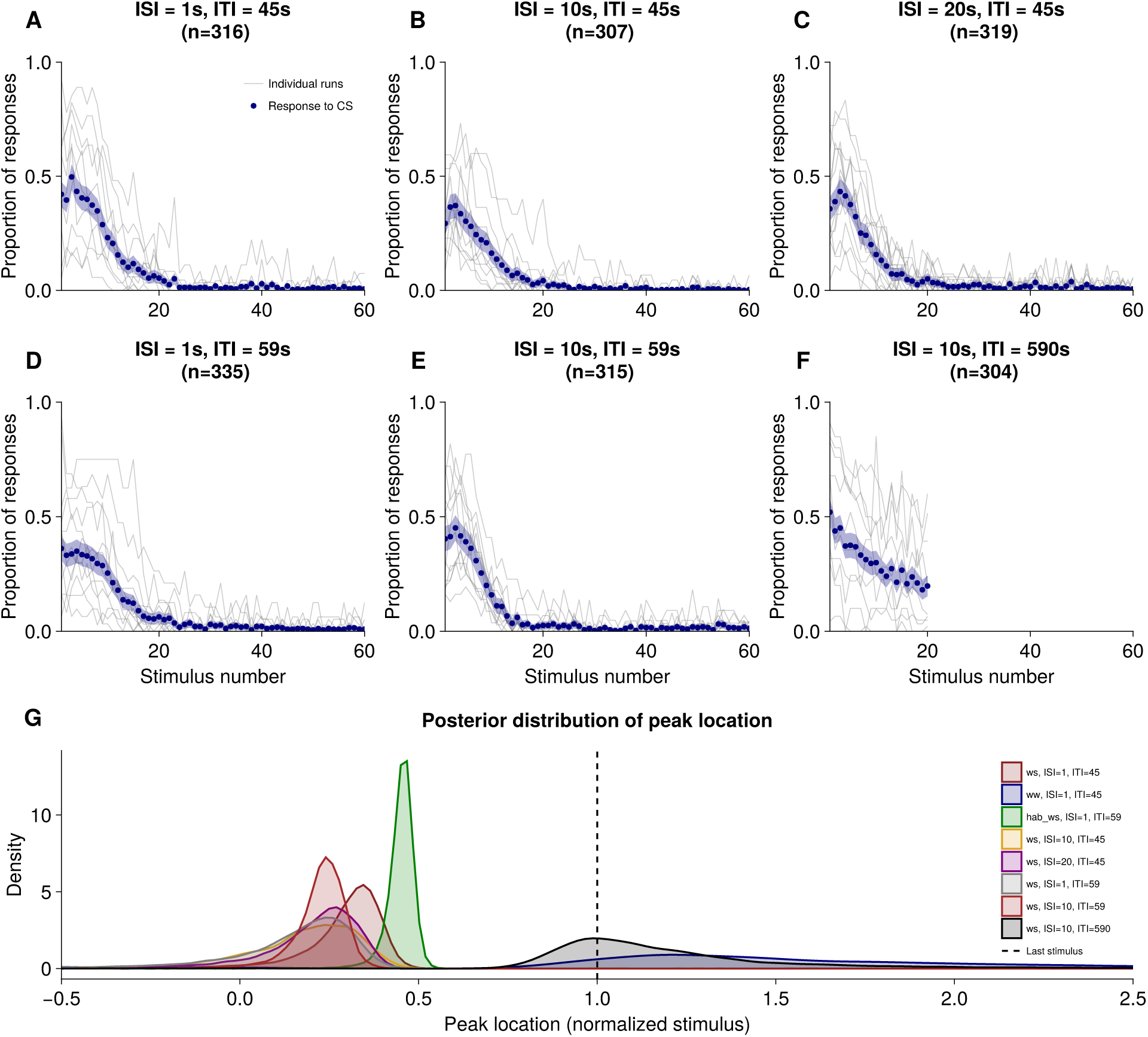
Temporal parameters modulate associative learning strength. Mean proportion contracted (navy points) and individual runs (gray lines) across conditions. Shaded regions represent 95% confidence interval (mean *±* 1.96 SEM; for proportions, 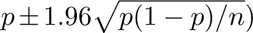. **(A)-(C)** ISI manipulation (ISI = 1, 10, 20s) at constant ITI = 45s. Panel **A** is the weak–strong condition of Figure 2B, replotted for comparison. **(D)-(F)** Combined ISI (1s and 10s) and ITI manipulation (59s and 590s). **(G)** Posterior distributions of the estimated peak location (*−β*_1_*/*(2*β*_2_)), in normalized stimulus units for each condition, derived from Bayesian quadratic regression. Each density represents the distribution across 20,000 posterior samples. The dashed vertical line marks the final stimulus (*T* = 10).

### 2.3 Learning criteria at single-cell level

As evident from Fig. 4, *Stentor* behavioral trajectories exhibit considerable variability across conditions and experimental runs. While population-level curves reveal non-monotonic responses to weak-strong pairing in most conditions, individual runs show substantial heterogeneity in both magnitude and timing of response enhancement. Such heterogeneity is characteristic of single-cell behavioral paradigms ^13,26^ but obscures key features of learning dynamics: the proportion of cells that learn, the speed of acquisition, and the strength of that learning at the individual cell level. To address these questions and establish a foundation for probing molecular mechanisms, we aim to identify individual cells that exhibit genuine associative learning—i.e., cells that acquire new responses to the weak tap based on its predictive relationship with the strong tap.

To illustrate associative learning at the single-cell level, consider two representative trajectories from the weak-strong protocol (Fig. 5A). One cell (magenta) exhibits the hallmark pattern of associative learning: initially non-responsive to weak taps (trial 1), it begins contracting during the pairing phase (trials 2-12), reaching a peak response rate around trial 8, before gradually habituating. This inverted-U pattern mirrors the population-level enhancement we reported above. In contrast, another cell (gray) shows monotonic habituation throughout training, never exhibiting enhanced responses despite identical stimulus exposure. These two examples help us build intuition for what drives the population-level enhancement: cells that initially fail to respond but gradually acquire contractions during pairing. Below, we characterize single-cell trajectories to capture these patterns across different conditions using a number of metrics that are robust to measurement noise, but sensitive enough to capture the behavioral trajectory we observed in the population response.

**Figure 5:**
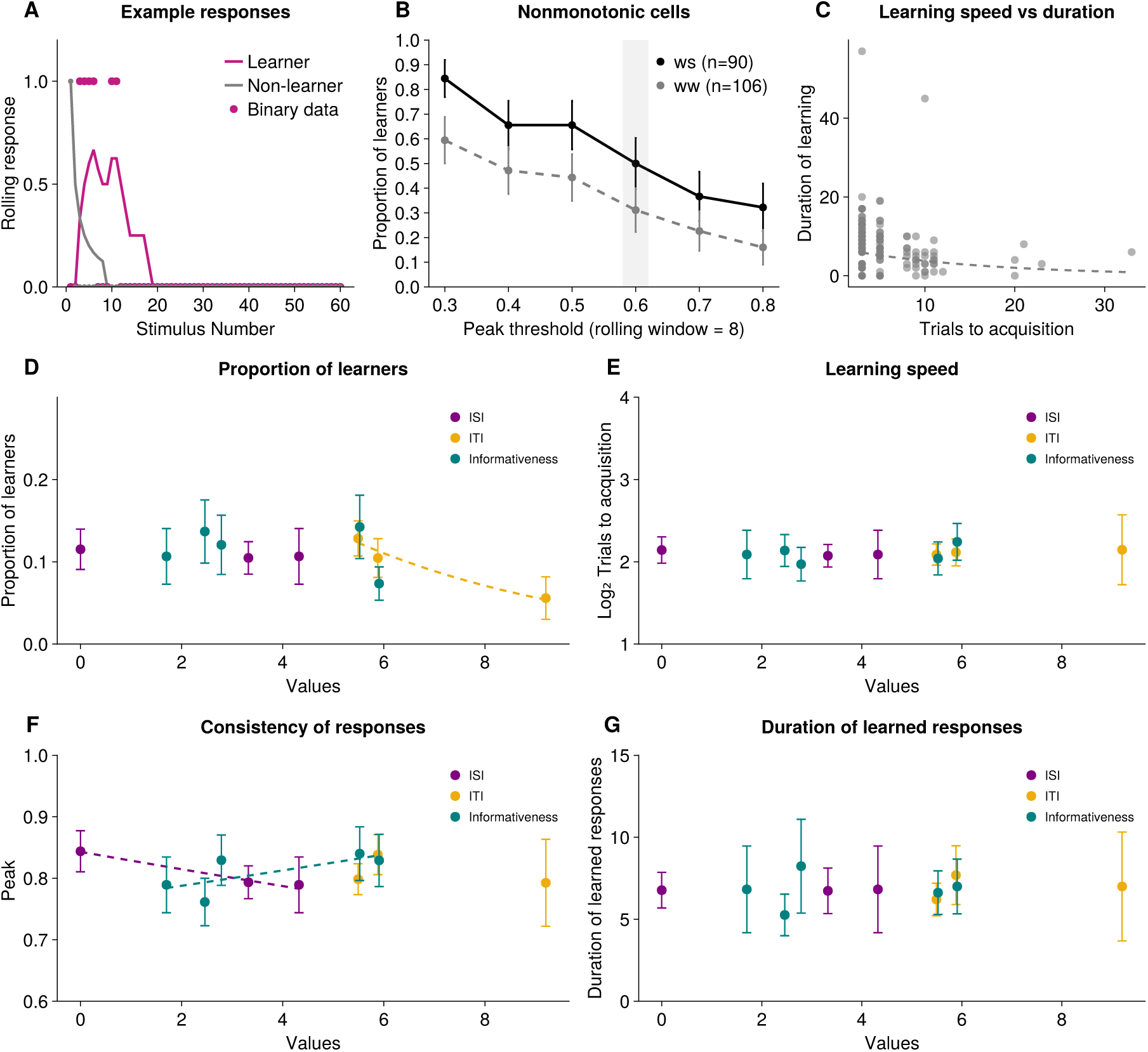
Characterization of associative learning at the single-cell level. **(A)** Sample responses illustrating learner versus non-learner profiles. Learner cell (magenta) initially shows no response to the conditioned stimulus, then acquires increasing contraction responses over successive trials before eventually habituating. Non-learner cell (gray) shows monotonic habituation throughout. **(B)** Proportion of non-monotonic cells meeting peak response thresholds (rolling window = 8 trials). Among cells exhibiting non-monotonic responses, weak-strong protocol yields a higher proportion meeting peak criteria than weak-weak control across all thresholds tested. Shaded region indicates the 60% threshold used for learner classification. Error bars indicate 95% confidence intervals, 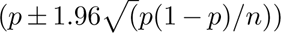 **(C)** Learning speed and duration of learned responses are negatively correlated. **(D)-(G)** Proportion of learners, speed of learning, consistency of responses and duration of learned responses as a function of inter-stimulus interval (ISI), inter-trial interval (ITI), and informativeness (ITI+ISI)/ISI. Dashed lines represent statistically significant relationships between independent and dependent variables. Learning speed and duration of learned responses are relatively insensitive to ISI, ITI and informativeness, but the proportion of learners declines significantly with longer ITI. The consistency of responses increases dramatically with shorter ISI.

Single-cell responses at each trial are binary (contraction or no contraction) and stochastic (e.g., due to membrane potential noise, among other factors), making individual trajectories noisy and difficult to interpret. To address this, we calculated rolling averages over a window of *k* trials, which smooth stochastic fluctuations while preserving response dynamics. We focused on cells that were initially non-responsive to weak taps but capable of responding. For each cell, we computed a rolling average of its contraction responses over the preceding 8 trials. We classified a cell as a learner if this average exceeded 60% for a sustained period during training. This metric ensures cells show learned responses consistently rather than sporadically. We characterized each learner by its *trials to acquisition* (when it first reached threshold, reflecting learning speed, cf. ^27^), *peak response* (the maximum of its smoothed trajectory, reflecting response consistency), and *duration of learned response* (how long it remained above threshold, reflecting persistence). All measures were robust to the choice of smoothing window and threshold (see Methods and Fig. S3).

Across all thresholds tested, weak-strong pairing consistently produced a higher proportion of learners than the weak-weak control (*β* = *−*3.60, *z* = *−*3.48, *p <* 0.001; Fig. 5B). As expected, increasing the threshold reduced the proportion of cells classified as learners (*β* = *−*12.49, *z* = *−*10.05, *p <* 10*^−^*^2^^3^). Importantly, the interaction between condition and threshold was not significant (*β* = 0.43, *z* = 0.25, *p* = 0.80). This result indicates that the two conditions decline at the same rate as the threshold increases, and that the proportion of learners in the weak-strong condition does not depend on a particular threshold choice.

### 2.4 ITI and ISI differentially control learning probability and strength

In this section, we investigated how temporal parameters shape different aspects of learning: whether cells learn, how quickly, how consistently, and for how long. We first tested whether temporal parameters predicted the proportion of learners using generalized linear models (details in Methods). ITI, the interval between one CS-US pairing and the next, was a significant predictor (*β* = -0.24, p = 0.001, Fig. 5D), with longer ITI associated with fewer learner cells. In contrast, ISI did not predict whether cells became learners (*β* = 0.0015, p = 0.97). Model comparison favored the model treating ITI and ISI as separate predictors over informativeness (the ITI/ISI ratio; log BF = -1.32, Fig. 5D), indicating that whether cells learn depends primarily on the inter-trial interval rather than overall predictive information.

We then tested whether temporal parameters predicted learning speed among cells that learned. Regression models revealed that neither ISI, ITI, nor informativeness reliably predicted trials to acquisition, and this result was robust across multiple model specifications (see Methods). This indicates that while ITI determines whether cells learn, once learning occurs, acquisition speed is relatively invariant across the temporal parameters we tested.

We next examined whether temporal parameters predicted learning strength among cells that learned. We found that ISI significantly predicts peak magnitude (*β* = -0.017, p = 0.012, Fig. 5F). Because peak reflects the maximum rolling average, this result suggests that cells trained with shorter inter-stimulus intervals responded more consistently after reaching acquisition, while those trained with a longer ISI showed more variable responding despite achieving acquisition. Model comparison favored the ISI alone model over the ISI + ITI model (log BF = -2.65) and informativeness (log BF = -0.666). ITI did not contribute to predicting peak magnitude.

Finally, we tested whether the duration of learned responses depended on temporal parameters. Consistent with our findings for trials to acquisition, neither ISI, ITI, nor informativeness reliably predicted persistence (all p *>* 0.05). Together, these results reveal that temporal parameters determine whether cells learn (ITI) and how strongly or consistently they show their learned responses (ISI), but not how quickly they acquire responses or how long they maintain them (see Fig. S4 for a breakdown of relationships).

### 2.5 Modeling associative learning in *Stentor coeruleus*

#### 2.5.1 Model setup

We developed a computational model of the non-monotonic learning dynamics we observed. The model formalizes the hypothesis that these dynamics emerge from two opposing processes: associative coupling between weak and strong stimuli, and differential habituation that causes both responses to decay with repetition.

Our model builds on empirically established features of *Stentor* ’s mechanosensory behavior, in particular intensity-dependent habituation, where responses to weak mechanical stimuli decay faster than responses to strong stimuli ^19,22^. We model each response as undergoing exponential decay at a stimulus-specific rate. Let *r_s_*(*n*) and *r_w_*(*n*) denote the response magnitudes to strong and weak stimuli on trial n, respectively. In the absence of pairing or when weak and strong stimuli are presented independently, these responses habituate according to:

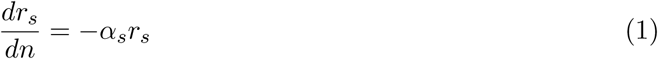

and

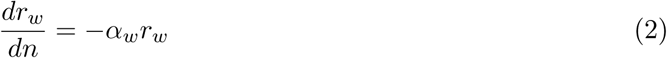

where *α_w_* and *α_s_* are habituation rate constants that determine how quickly responses decay, with a larger rate constant corresponding to faster habituation. Because weak stimuli produce faster habituation than strong stimuli, we impose the constraint *α_w_ > α_s_*.

For the weak-strong pairing protocol, where weak stimuli reliably precede strong stimuli, we hypothesize that weak stimuli acquire anticipatory responses to the upcoming strong stimulus. This is reflected in the increased contractions to weak taps we observed during early training. We formalize this through an associative coupling term *α_c_* that captures how response strength from the strong stimulus *r_s_* transfers to the weak stimulus *r_w_*. When *α_c_* = 0 (either because there is no pairing or because the coupling is absent), the stimuli are independent and both undergo simple habituation (Equations 1-2). When *α_c_ >* 0, the strong response drives the weak response, counteracting habituation and producing elevated weak tap responses. This yields the coupled dynamics:

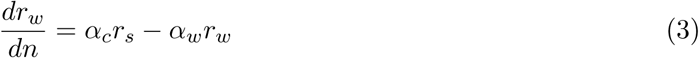

where the first term represents associative drive and the second term represents habituation (Fig. 6A). Solving this differential equation yields:

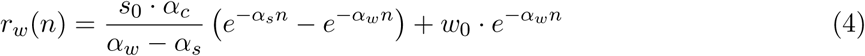

**Figure 6:**
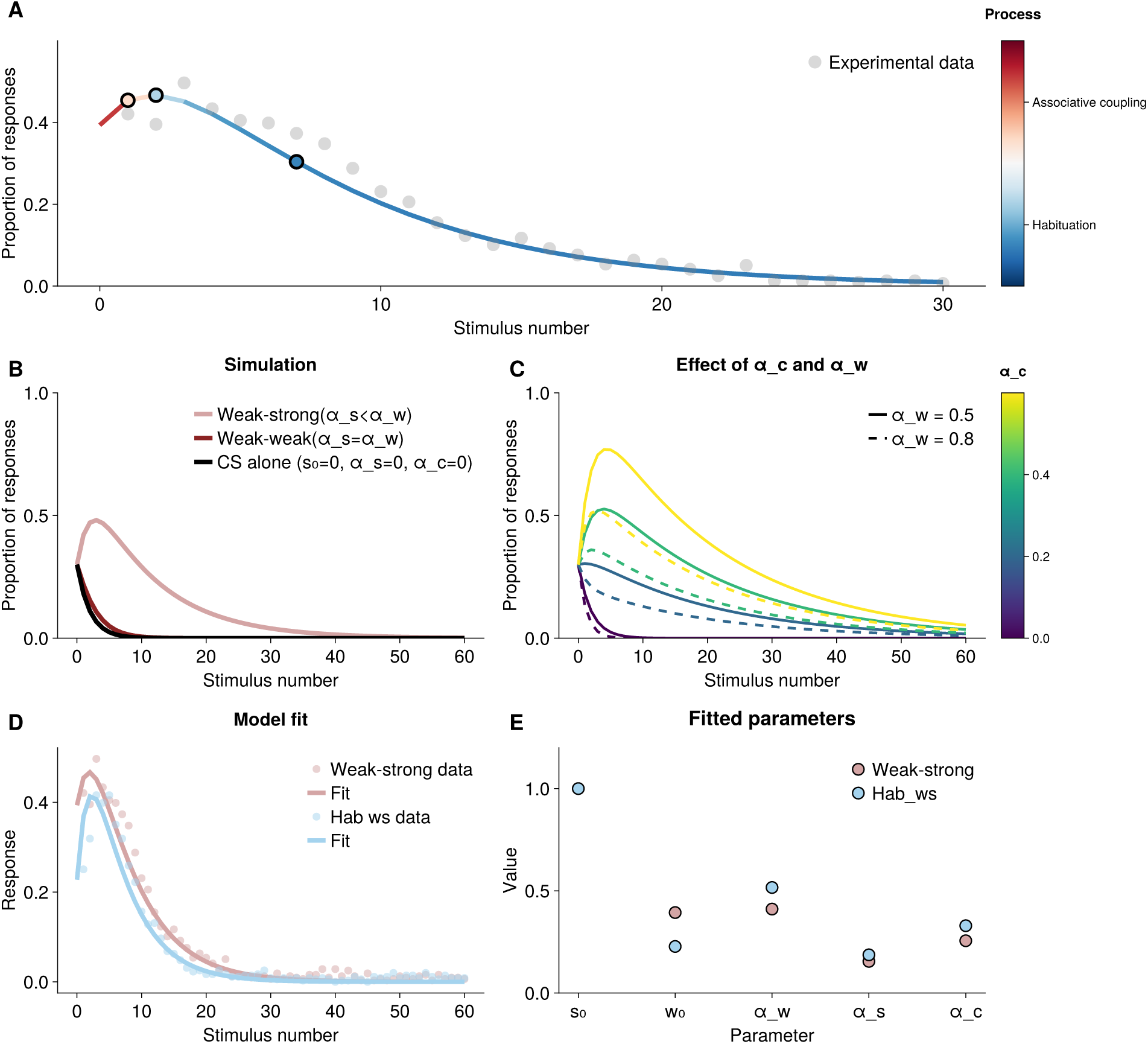
Associative learning model captures non-monotonic learning dynamics through competition between associative coupling and habituation. **(A)** Illustration of the competition between associative drive (*α_c_r_s_*) and habituation (*α_w_r_w_*) over time. Solid curve shows the net response *r_w_*(*n*), decomposed into contributions from associative drive (red region, early trials) and habituation (blue region, later trials). Gray points: empirical data from weak-strong pairing (ISI = 1 s, ITI = 45 s, same data as in Fig. 2B). note the transient rise and fall of the population responses, consistent with experimental observations. **(B)** Model simulations across pairing conditions. Weak-strong pairing (pink; *α_s_ < α_w_*) produces non-monotonic dynamics. Weak-weak pairing (dark red; *α_s_* = *α_w_*) and CS alone (black; *s*_0_ = 0, *α_c_* = 0) produce monotonic habituation. **(C)** Parameter sensitivity analysis. Coupling strength (*α_c_*, color gradient) controls peak magnitude. Habituation rate (*α_w_*, solid vs. dashed lines) controls rise and fall dynamics. Fixed parameters: *s*_0_ = 0.8, *w*_0_ = 0.3, *α_s_* = 0.05. **(D)** Model fits to empirical data. Conditions used data from weak-strong pairing ISI = 1s, ITI = 45s and habituation (ISI =1s), and weak-strong pairing ISI=1, ITI=59 (same data as in Fig. 2B and G, respectively) The model captures both non-monotonic patterns. **(E)** Comparison of parameters extracted from model fits in weak-strong and habituation weak-strong conditions.

where *s*_0_ and *w*_0_ are the initial response magnitudes to strong and weak stimuli, respectively. The competition between two processes produces non-monotonic dynamics (Fig. 6A). Early in training, *r_s_* is large, so the associative term dominates and increases the response to the weak stimulus (red phase). As both responses habituate over trials, *r_s_* declines, weakening the associative drive. Eventually, direct habituation overtakes the diminishing drive from *r_s_*, causing the response to the weak stimulus to decline (blue). The model thus produces an inverted-U trajectory consistent with our empirical observations.

#### 2.5.2 Model simulations and validation

We validated our model through simulations and fits to empirical data using parameters estimated from strong-tap habituation (*s*_0_ = 0.8*, α_s_* = 0.05) and baseline weak-tap responses (*w*_0_ = 0.3; more fitting details in Methods). The model correctly predicted qualitatively different outcomes across pairing conditions (Fig. 6B). Weak-strong pairing produced robust non-monotonic dynamics, while weak-weak pairing produced monotonic habituation. We systematically varied coupling strength (*α_c_*) and habituation rate (*α_w_*) (Fig. 6C). Increasing *α_c_* amplified peak responses, while increasing *α_w_*accelerated both the rise and the fall.

We fit the model to weak-strong and habituation weak-strong data (Fig. 6D,E). For both conditions, the model was able to capture the early non-monotonic pattern and the fast decline.

## 3 Discussion

Our study broadens the scope of learning capabilities in protozoa by providing evidence for associative learning in *Stentor coeruleus*. Temporally pairing weak and strong mechanical stimuli (taps) induced a transient enhancement of responding (contraction) to the weak tap. This phenomenon appears to depend strongly on the temporal pairing protocol; it is absent in a variety of other protocols involving weak and strong taps but lacking consistent temporal pairing. Two features of our data are nonetheless compatible with non-associative sensitization account, namely a transient enhancement and fewer learners at longer ITIs. We showed with a common test procedure that *Stentor* were able to maintain their memory of the pairing, whereas cells that received the same strong taps without pairing did not.

Our protocol is rather unusual in using the same stimulus (with different intensities) for the CS and US. Typically, the CS and US are drawn from different sensory modalities and elicit qualitatively different responses. Because the response repertoire of *Stentor* is fairly limited, we made this design choice in order to build on past results using these stimuli in mechanosensory habituation paradigms ^19,22^. While unconventional, our choice is not without precedent: a fear conditioning study in rats, in which weak and strong electric shocks were paired, also produced a non-monotonic learning curve ^23^. Future work could incorporate additional sensory modalities, such as light stimuli ^28,29^, to test conditioning in *Stentor*.

Pavlovian conditioning in animals often obeys a strong quantitative law: the number of conditioning trials until an acquisition criterion is met depends logarithmically on the ITI/ISI ratio, also known as *informativeness* ^30^. This law implies a form of timescale invariance: rescaling the ISI and ITI by a constant should not affect the trials to acquisition, consistent with the findings from many studies. To test the applicability of this law to conditioning in *Stentor*, we parametrically manipulated the ISI and ITI. We found that neither parameter (nor their ratio) correlated with trials to acquisition (defined as the peak of the learning curve), but ITI negatively correlated with the proportion of learners, and the ISI negatively correlated with response consistency among learners. These results suggest that the quantitative laws of conditioning in animals do not straightforwardly apply to *Stentor*, though two important caveats should be borne in mind. First, the non-monotonic structure of the learning curve in our study makes challenging a direct comparison with monotonic learning curves in animals. Second, the long latency of re-extension following a contraction limited the range of feasible temporal parameters.

A simple mathematical model, inspired by the Rescorla-Wagner model^31^, could explain the non-monotonic structure of the learning curves we observed. The model combines two processes: (i) an associative learning process by which the weak tap acquires a predictive relationship with the strong tap, and (ii) a habituation process by which the responses to the strong and weak taps both decay with repetition. The associative learning process produces the enhancement, which is made transient by the habituation process. This model is primarily a proof of concept; we don’t claim that it is a comprehensive explanation of our data, and indeed it cannot explain the quantitative dependence on ITI and ISI parameters. Ultimately, we hope to replace this abstract model with a more mechanistically detailed one that can be linked to the molecular biology of learning in *Stentor*. Some promising steps in this direction have already been made for modeling habituation ^20,32^.

Our findings have broad implications for the evolutionary origins of associative learning. If ciliates such as *Stentor* and *Paramecium* are capable of associative learning (see ^11^ for a review of past studies), then there must be a cellular mechanism for such learning that does not depend on synaptic modification. This compels us to consider fundamentally different mechanisms. It has been suggested that such mechanisms might also operate in the brain^33,34^—a conjecture that invites a new program of research bridging the study of associative learning in protozoa and metazoa.

## 4 Methods

### 4.1 Code and data availability

All data and analysis code is provided in a GitHub repository: https://github.com/nhithuydoan/associative_learning_Stentor

### 4.2 Cultures and maintenance

*Stentor coeruleus* cells were obtained from Carolina Biological (131598) and grown in filtered spring water (Carolina Biological, 132450) for at least two weeks prior to experiments. Cultures were kept in Pyrex bowls in dark drawers at 22°C when not in use.

*Chlamydomonas reinhardtii* (Chlamy Center, CC-125) were used as *Stentor* ’s food source. *Chlamydomonas* were grown on 1.5% agar in TAP media (Phytotechlabs, T8224) under continuous bright light. For feeding preparations, *Chlamydomonas* cells were transferred from agar plates to 50 mL of filtered spring water using five loopfuls of a sterile inoculating loop. Large fragments of *Chlamydomonas* were broken up with gentle pipetting and shaking. *Stentor* cultures (500 mL) were fed with 7 mL of feeding solution every four to five days.

Cultures of *Stentor* and *Chlamydomonas* were inspected weekly for signs of mold, dead cells, or slow growth. *Stentor* were transferred to new cultures with fresh water each month to maintain health and avoid bacterial overgrowth.

### 4.3 Experimental paradigm

The experimental apparatus included a large, dark, custom-built acrylic box mounted on an isolation system (ThorLabs, PTT600600). Four sorbothane feet attached to a breadboard (ThorLabs, AV6) provided additional vibration isolation. Three custom experimental devices were cut from 1/8” thick acrylic sheet and evenly spaced on the breadboard. The solenoids (Adafruit Mini Push-Pull 5V) were placed in the center of each device, connected to a programmable power supply (KORAD KA6003P) and controlled by a Raspberry Pi. When energized, the solenoid piston moved up 2.5 mm, contacted the dishes, and returned to their original position.

For image capture, each device was equipped with a mounted camera (Edmund Optics, BFS-U3-120S4M-CS) and a 24-bit LED ring. The LED ring was covered with diffusion paper to reduce glare and provide uniform illumination. The cameras recorded experiments at 10 FPS. Cell behaviors were captured 1 second before and 0.9 seconds after stimulus onset, so that the measurement window closes before the strong tap in the weak–strong protocols. Unless otherwise noted, contraction score is defined as the comparison of these two windows for the weak stimulus, and the same window is used in all conditions regardless of ISI. Rolling averages are computed from these scores.

### 4.4 Experimental preparation

For each experiment, 400 *µ*L of cultured *Stentor* were transferred to 2300 *µ*L of filtered water in a 35 mm Petri dish coated with poly-D-lysine (Mattek). Cells were left undisturbed for at least eight hours and exposed to dim light (3 fc) for two hours prior to the experiment. Experiments were carried out across different times of day with replicates from multiple independently cultured populations. All experiments were carried out at 22°C by a single experimenter.

Unless otherwise noted, all behavioral protocols (Fig. 1D) deliver 60 mechanical stimuli with different inter-trial interval (ITI) and inter-stimulus interval (ISI).

### 4.5 Measurement of behaviors

*Stentor* has the tendency to detach and swim away, aggregate in groups, and intermittently undergo mitosis or conjugation. These behaviors create challenges because we cannot ensure cells receive the same number of mechanical taps or stimulus intensity, nor can we rule out whether reorganization of nuclear material occurs during conjugation (sexual reproduction) or mitosis changes alter their mechanosensory responses. Responses for grouped *Stentor* are ambiguous because grouped cells experience mechanical/hydrodynamic interference between neighbors ^21,35^, making it unclear whether contractions result from the delivered stimulus or from neighboring activity. For these reasons, we excluded cells that underwent mitosis or conjugation, swam away during one or more stimuli, or contacted neighboring cells. Only cells that remained anchored throughout the experiment, maintained trumpet-shaped morphology, and showed minimal contact with neighbors were selected for analysis.

Behavioral data were pre-processed using a custom Julia script where researchers manually selected cells according to the listed criteria and scored contraction responses. A cell was considered contracted if it curled in a sphere-like shape (Fig. 1B). If the cell was contracted before a stimulus (spontaneous contraction), we mark as NA and do not analyze it. The 45-s inter-trial interval was chosen so that cells return to a fully extended baseline before the next trial begins. On rare occasions a cell had not fully re-extended before the next stimulus onset; that trial was excluded for that cell, which occurred in roughly 0.1% of trials. Between researchers, the contraction scores shared 93% similarity.

### 4.6 Statistical analyses

#### Experimental design and sample sizes

Experimental conditions were tested across 10-20 independent runs, with each run yielding 10-40 cells for a total of approximately 300 cells per condition. Sample sizes (n) reported throughout represent total cells aggregated across runs. Besides inter-stimulus interval (the interval between CS and US or weak tap and strong taps) and inter-trial intervals (the interval between one pair of CS-US to the next pair), we defined informativeness as the ratio (ISI + ITI)/ISI. All temporal variables (ISI, ITI, informativeness) were *log*_2_-transformed prior to analysis.

#### Population-level analyses

To quantify non-monotonic response patterns at the population level, we used mixed-effects logistic regression with stimulus number and its square as fixed effects and random intercepts for experimental runs to account for variability within runs. In addition, we fit Bayesian quadratic regressions for each condition, with stimulus values normalized to [0, 1]. Because response curves frequently exhibit a long flat tail beyond the peak, fitting the full sequence would allow this tail to dominate and obscure the non-monotonic structure of interest; we therefore restricted the regression to the first 10 stimuli. The model took the form:

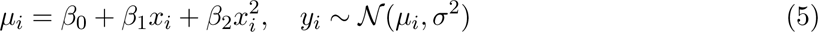

with priors *β*_0_ *∼ N* (0.4, 0.3), *β*_1_ *∼ N* (0, 1), *β*_2_ *∼ N* (0, 1), and *σ ∼ N* ^+^(0, 0.2). Posterior inference was performed using the No-U-Turn Sampler with 20,000 samples, implemented in Turing.jl^36^. From the posterior, we computed the probability that the quadratic was concave (*β*_2_ *<* 0) and that its peak, located at *−β*_1_*/*(2*β*_2_), fell within the normalized stimulus range (0, 1), where 1 corresponds to the final stimulus (*T* = 10). This gives a direct estimate of the probability that the response peaked before the end of the stimulus sequence (Fig. 4G).

For control comparisons, we used one-sided paired *t*-tests. Shaded areas and errors bar represent 95% confident intervals (mean *±* 1.96 SEM; for proportions, 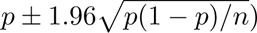.

#### Quantifying individual response trajectories

We restricted analysis to cells capable of demonstrating acquisition of a new response. Cells that contracted on the first weak tap were already responsive before training and could not demonstrate learning. Cells that never contracted throughout the experiment provided no information about learning capacity. We therefore analyzed only cells that were initially non-responsive to weak taps but showed at least one contraction during the experiment. Of all cells, 31.5% never responded to the weak taps.

To classify each cell as a *learner* or *non-learner*, we first converted the binary contraction responses into a rolling contraction probability. For each trial, we computed a rolling average by averaging the responses the preceding *k* trials, where *k* is a window size. Window size involves a tradeoff: short windows (*k* = 3 trials) are sensitive to spurious peaks from random responses, while long windows (*k* = 12 trials) oversmooth the signal and obscure response dynamics. To select an appropriate value, we asked which *k* best distinguished the weak–strong condition from the weak–weak control, which reflects the non-monotonic and habituation profiles at population-level patterns. Windows of *k* = 5 and *k* = 8 both satisfied this criterion and produced quantitatively similar learner classifications (mixed-effects logistic regression: *β* = *−*0.39, *z* = *−*1.70, *p* = 0.090; Fig. S3). Because *k* = 8 is stricter and thus more conservative in classifying cells as learner, we report results using *k* = 8 throughout, though all statistical findings hold with *k* = 5.

#### Learner classification and threshold validation

A cell was classified as a learner if its smoothed response trajectory exceeded a threshold level of consistent responding. We tested thresholds from 30% to 80% and validated this classification using a mixed-effects logistic regression (Bernoulli GLM with logit link) predicting learner status from condition (weak–strong vs. weak– weak), threshold, and their interaction, with a random intercept for each cell. We selected a 60% threshold to balance sensitivity against false positives while maintaining sufficient sample size for subsequent analyses. This choice is supported by the non-significant condition *×* threshold interaction reported in the Results, which indicates that the proportion of learners in the weak– strong condition does not depend on a particular threshold choice.

#### Learning metrics

We quantified three aspects of single-cell learning dynamics. *Trials to acquisition* is the first trial at which a cell’s *k*-trial rolling average exceeds the 60% threshold, which we use as a measure of learning speed. *Peak response* is the maximum value of the rolling average during training, which measures response consistency. *Duration of learned response* is the number of trials between a cell first crossing above and last falling below the 60% threshold, serving as a measure of learning persistence. These last two metrics are particularly relevant for *Stentor* because mechanosensory stimuli naturally induce habituation; characterizing both strength and persistence provides a more complete picture of associative learning dynamics in this system.

#### Single-cell learning analyses

We fit two generalized linear models: Model 1 with informativeness as a single predictor, and Model 2 with ISI and ITI as separate predictors. Distributions were chosen based on the nature of each dependent variable: binomial family with logit link for learner status (binary); Gamma family with log link for peak magnitude. For trials to acquisition and duration of learning, we tested multiple distribution families (normal and Gamma for trials to acquisition; Poisson and Gamma for duration of learning) to ensure robustness of findings. For trials to acquisition specifically, we also fit Model 3 (mixed-effects model with random intercepts for each cell) and Model 4 (Gamma distribution with log link). All models and distribution choices yielded qualitatively similar results, with none of the predictors showing significant relationships with trials to acquisition or duration of learning.

Models were compared using the Bayesian Information Criterion (BIC), with log Bayes factors approximated as log BF *≈* 0.5ΔBIC, where ΔBIC is the difference between the BIC for two models.

### 4.7 Model of associative learning

#### Model specification

The associative learning model (Equations 3–4) has five parameters: *s*_0_ (initial strong response), *w*_0_ (initial weak response), *α_s_* (strong habituation rate constant), *α_w_*(weak habituation rate constant), and *α_c_* (coupling strength). To enforce the constraint *α_w_ > α_s_*, we reparametrized *α_w_* = *α_s_* + *δ* + *α_c_*, where *δ >* 0 represents the additional habituation rate beyond the strong-tap baseline. We included *α_c_* in this reparametrization rather than setting *α_w_* = *α_s_* + *δ* alone, as this provided better fits to the data. This formulation implies that coupling strength contributes to the effective habituation rate of the weak response, with a stronger associative link facilitating faster decay.

#### Model simulations

We simulated three scenarios to examine how model parameters shape learning dynamics. All simulations ran for 60 trials. In the weak-strong condition, we set *s*_0_ = 0.8, *w*_0_ = 0.3, *α_s_* = 0.1, *α_w_* = 0.5, *α_c_* = 0.4 to produce non-monotonic dynamics through strong associative drive. In the weak-weak condition, we set *s*_0_ = *w*_0_ = 0.3, *α_s_*= *α_w_*= 0.5, *α_c_* = 0.2 to simulate the case where both stimuli habituate at equal rates. In the CS-alone condition, we set *s*_0_ = 0, *α_s_* = 0, and *α_c_* = 0, eliminating both the US response and associative coupling, producing simple exponential habituation.

#### Parameter estimation

We fit the model to population-averaged response data separately for weak-strong condition using nonlinear least squares (Levenberg-Marquardt algorithm; LsqFit.jl). We first fit strong-tap-only data (ISI = 1s, ITI = 45s) to exponential habituation (*r_s_*(*n*) = *s*_0_*·e^−α_s_n^*) to obtain initial estimates of *s*_0_ and *α_s_*. We then fit weak-tap responses to Equation 4, using these estimates as initial values for *s*_0_ and *α_s_*, and simultaneously estimating all five parameters (*s*_0_, *w*_0_, *δ*, *α_s_*, *α_c_*). Initial values were: *s*_0_ and *α_s_* from stage 1, *w*_0_ set to the first trial response, *δ* = 0.01, and *α_c_* = 0.5. Parameter bounds were: *s*_0_ ∈ [0.7, 1.0], *w*_0_ ∈ [0.01*, ∞*), *δ* ∈ [0.0001*, ∞*), *α_s_* ∈ [0.0001*, ∞*), *α_c_* ∈ [0.05*, ∞*).

All analyses and visualizations were performed in Julia version 1.12.

## 5 Acknowledgments

The work was supported by the Air Force Office of Scientific Research grant (FA9550-22-1-0345), a grant from the Alfred P. Sloan Foundation through the Matter-to-Life Program (2026-79563), and a Schmidt Sciences Polymath Award. We thank Matt Laudon for advice on our *Chlamydomonas* culture, Ed Soucy and Yuwei Li for help with our experimental apparatus, Peng Qian, Shuze Liu, Arthur Prat-Carrabin, and Jake Upton for feedback on earlier drafts. Zach Kelso, Yue Miao, John Vastola, and Maddie Snyder provided helpful discussion throughout the project.

## 6 Supplemental figures

**Figure S1:**
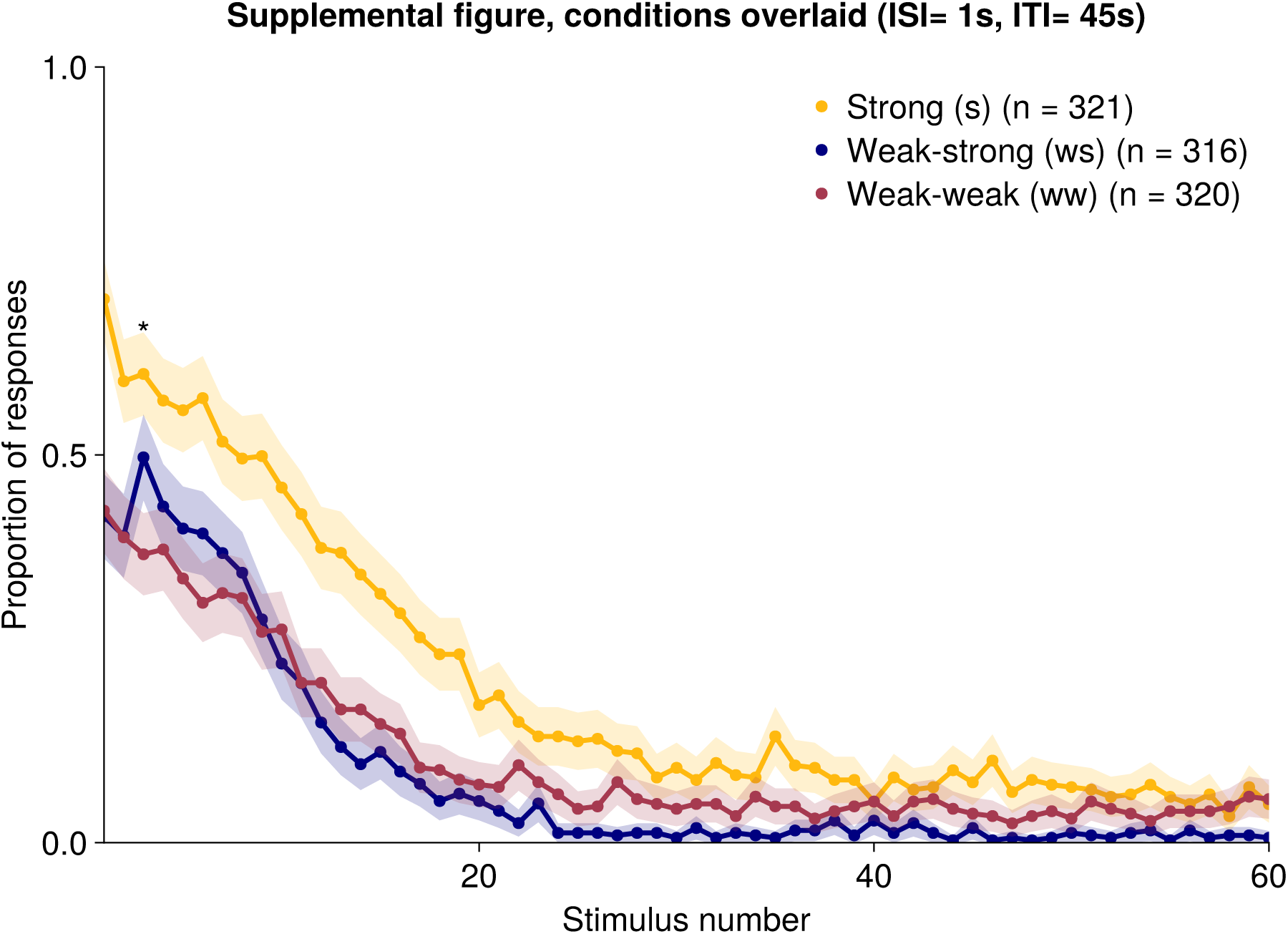
Comparison between strong, weak–strong, and weak–weak conditions. Strong alone (s; yellow; *n* = 321 cells, 19 runs), weak–strong (ws; navy; *n* = 316 cells, 10 runs), and weak–weak (ww; dark red; *n* = 320 cells, 14 runs). Points are the mean proportion of cells contracting at each stimulus. Shaded areas are 95% confidence intervals for proportions 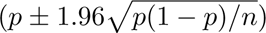. Same data as Fig. 2A–C. Asterisk denotes significance at stimulus 3 between weak–strong and weak-weak conditions (Welch *t*(14.2) = 1.83, one-sided *p* = 0.044).

**Figure S2:**
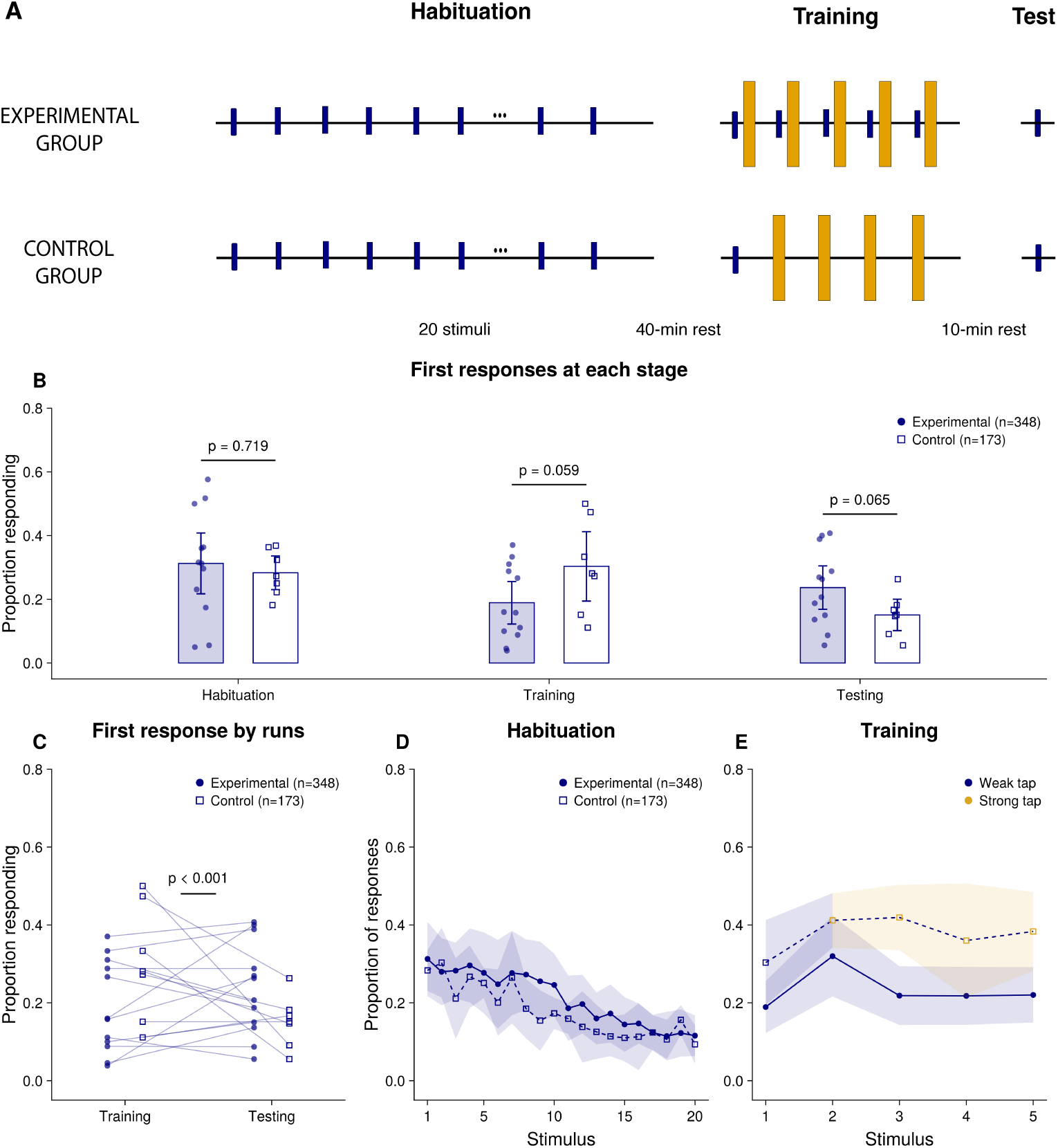
Common test after different training protocols. **(A)** Experimental design for Set 2 data. Similar to design for Set 1 data (Fig. 3), both groups receive identical pre-habituation (20 weak taps) and an identical test (one weak tap), and differ only during training: five weak– strong pairings (experimental) versus one weak tap followed by four strong taps (control). Rests were 40 min before training and 10 min before test. Navy lines/dots are weak taps; yellow ones are strong taps. Filled navy circles represent experimental condition (*n* = 348, 12 runs); open squares are control (*n* = 173, 7 runs).**(B)** Responses to the first weak tap of each phase. Bars are means across runs, points are individual experimental runs, error bars are 95% CIs. Each *p* is for condition (response *∼* condition + (1*|*run), fit per phase). **(C)** The same first-tap responses at training and test, one line per run; *p* is the condition*×*phase interaction p-value. **(D)** Pre-habituation curve; points are means across runs; shaded area reprepresents the 95% CI. **(E)** Training curve, as in (D). All models are mixed-effects logistic regressions (Bernoulli, logit link).

**Figure S3:**
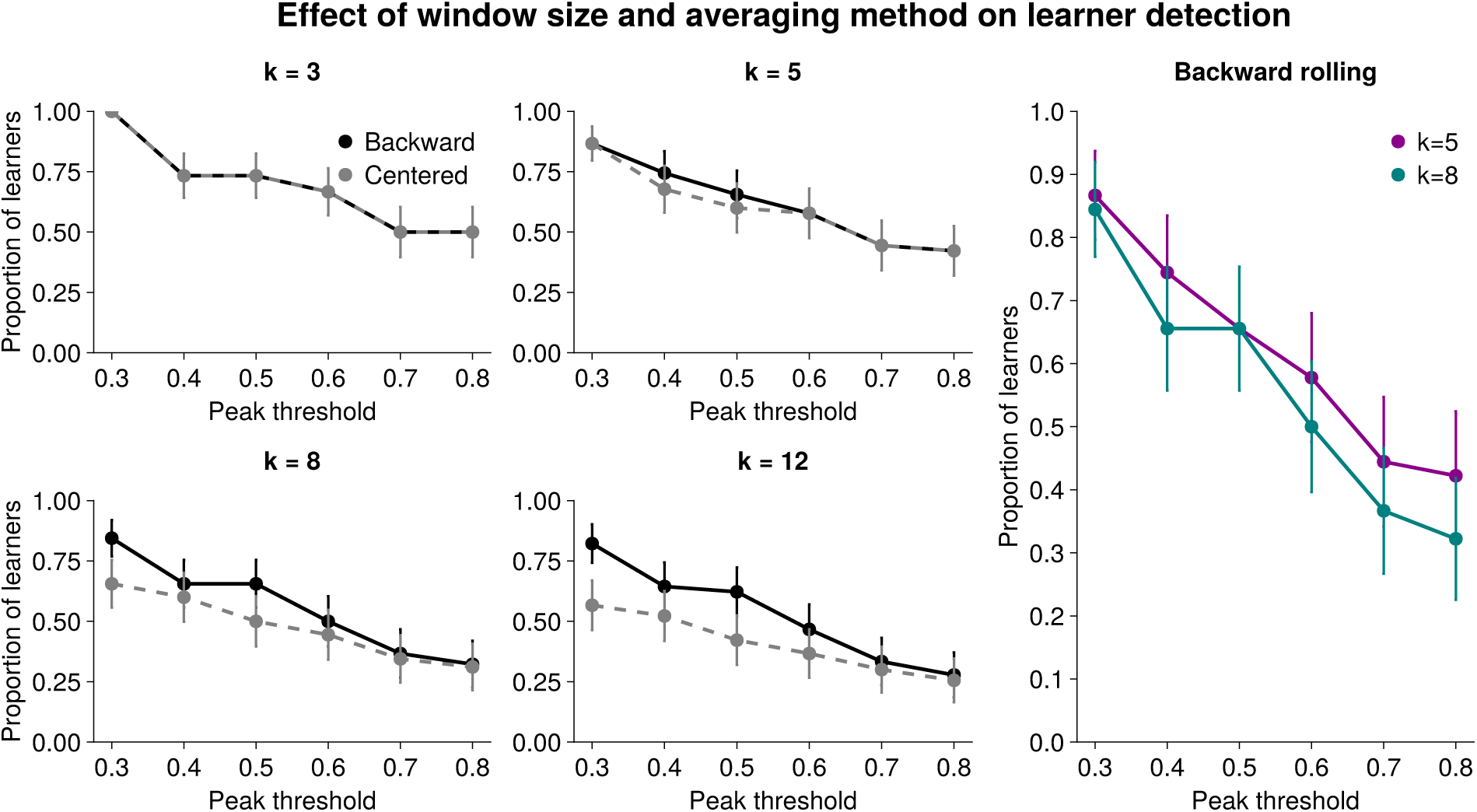
Learner classification is robust to smoothing parameter choices. Proportion of non-monotonic cells classified as learners across peak thresholds (30–80%) using backward and centered rolling averages with window sizes *k* ∈ {3, 5, 8, 12}. Error bars represent 95% confidence intervals. Backward and centered averaging produce similar results at small window sizes, though the two methods diverge as *k* increases. Our preferred windows of *k* = 5 and *k* = 8 yield quantitatively similar learner classifications (rightmost panel), with *k* = 8 producing slightly fewer learners on average (mixed-effects logistic regression: *β* = *−*0.39, *z* = *−*1.70, *p* = 0.090). We report results using *k* = 8 throughout as the more conservative estimate, even though all statistical findings hold with *k* = 5.

**Figure S4:**
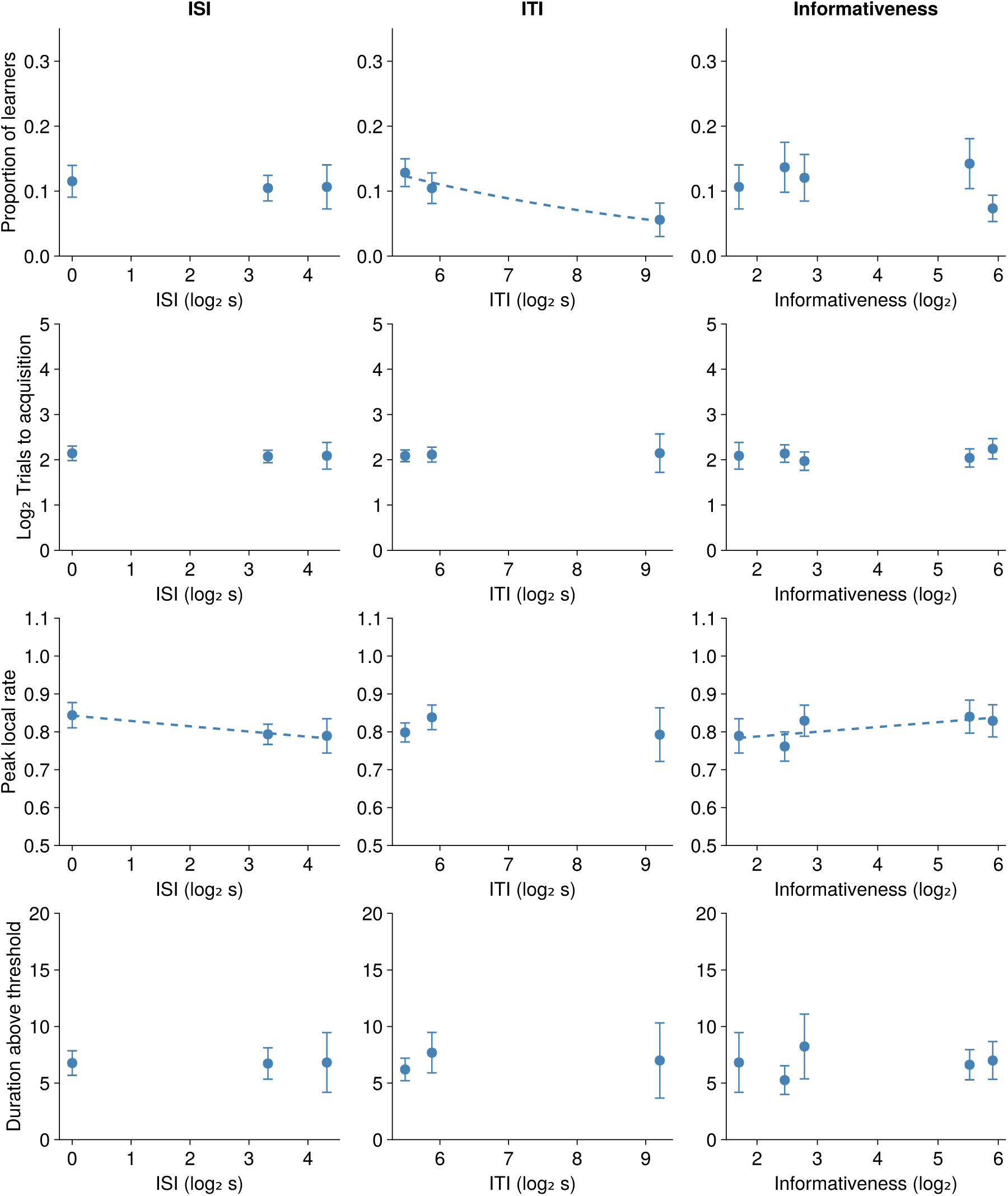
Temporal parameters differentially influence associative learning dynamics. Gray dashed line indicates significant regression relationships (*p <* 0.05). Proportion of learners (first row), trials to acquisition (second row), peak height (third row), and duration of learned response (fourth row) as a function of ISI, ITI, and informativeness (first, second, and third row respectively). Learning probability decreases significantly with longer ITI but shows no significant relationship with ISI or informativeness. Peak local rate decreases with longer ISI and lower informativeness (gray dashed line) but shows no relationship with ITI.

